# Social status alters chromatin accessibility and the gene regulatory response to glucocorticoid stimulation in rhesus macaques

**DOI:** 10.1101/365049

**Authors:** Noah Snyder-Mackler, Joaquín Sanz, Jordan N. Kohn, Tawni N. Voyles, Roger Pique-Regi, Mark E. Wilson, Luis B. Barreiro, Jenny Tung

**Author notes:** Correspondence to: Jenny Tung and Noah Snyder-Mackler.

## Abstract

Low social status is an important predictor of disease susceptibility and mortality risk in humans and other social mammals. These effects are thought to stem in part from dysregulation of the glucocorticoid (GC)-mediated stress response. However, the molecular mechanisms that connect low social status and GC dysregulation to downstream health outcomes remain elusive. Here, we used an *in vitro* glucocorticoid challenge to investigate the consequences of experimentally manipulated social status (i.e., dominance rank) for immune cell gene regulation in female rhesus macaques, using paired control and GC-treated peripheral blood mononuclear cell samples. We show that social status not only influences immune cell gene expression, but also chromatin accessibility at hundreds of regions in the genome. Social status effects on gene expression were less pronounced following GC treatment than under control conditions. In contrast, social status effects on chromatin accessibility were stable across conditions, resulting in an attenuated relationship between social status, chromatin accessibility, and gene expression post-GC exposure. Regions that were more accessible in high status animals and regions that become more accessible following GC treatment were enriched for a highly concordant set of transcription factor binding motifs, including motifs for the glucocorticoid receptor co-factor AP-1. Together, our findings support the hypothesis that social status alters the dynamics of GC-mediated gene regulation, and identify chromatin accessibility as a mechanism involved in social stress-driven GC resistance. More broadly, they emphasize the context-dependent nature of social status effects on gene regulation and implicate epigenetic remodeling of chromatin accessibility as a contributing factor.

## INTRODUCTION

Social status is an important predictor of health and survival in social animals, including humans (1-3). In the United States, for instance, men in the lowest income percentile live 15 year shorter lifespans, on average, than men in the highest income percentile (4). Similarly, British civil servants in the lowest employment grades are twice as likely to develop cardiovascular disease compared to those in the highest employment grades, despite universal access to the British National Health Service (5). Social status gradients in health can be explained in part by correlated variation in health risk behaviors, resource access, and the long-term effects of social adversity earlier in life (6, 7). However, increasing evidence, particularly from animal models, indicates that low social status also leads to direct physiological effects (8-13). For example, experimentally induced low social status in cynomolgus macaques and rhesus macaques leads to increased rates of atherosclerosis and hyperinsulinemia, and an exaggerated gene regulatory response to bacterial lipopolysaccharide (LPS) (8, 9, 14-21).

The direct effects of low status may be mediated, at least in part, by the chronic stress of social subordination (22, 23). In humans and other hierarchically organized primates, low status is associated with greater social unpredictability and reduced social control, fewer affiliative interactions, and reduced social integration (24). These experiences are thought to increase exposure to stressors (e.g., via increased rates of harassment) and reduce the capacity to buffer against stressful experiences (e.g., via lack of social support) (24). Such changes in turn lead to physiological alterations in the regulation of the hypothalamic-pituitary-adrenal (HPA) axis, which controls the production of glucocorticoids (GC), a class of steroid hormones involved in metabolic and immune regulation (25). Chronic social stress has been associated with both elevated baseline GC levels (e.g., (26, 27); but see (28)) and impaired GC negative feedback (i.e., GC resistance; (23, 29)), suggesting a physiological basis for social gradients in metabolic and inflammation-related disorders (30). In support of this hypothesis, the psychological stress of caring for a chronically ill child is associated with both GC resistance and impaired GC suppression of proinflammatory signaling (29). Similarly, major social stressors such as job loss or bereavement predict both GC resistance and susceptibility to the common cold (23). Causal effects of social stress on GC function have also been demonstrated in experimental animal models. For instance, red squirrel mothers exposed to cues for increased resource competition (but not increased resource competition itself) exhibit elevated GC levels, a change that in turn alters offspring growth (31).

At the molecular level, GCs operate as signaling molecules that affect transcriptional regulation, thus linking multi-organ HPA axis regulation to changes in metabolic and immune function within cells. Circulating GCs canonically influence gene regulation by diffusing across the cell membrane and binding to the cytosolic form of the nuclear receptor glucocorticoid receptor (GR, encoded by the gene *NR3C1*). Once bound, the GC-GR complex translocates into the nucleus where it binds directly or indirectly (via tethering to other transcription factors: (32) to target DNA sequences (e.g., glucocorticoid response elements, GREs; (25, 33)). There are multiple potential mechanisms through which GCs exert anti-inflammatory effects, and GCs have been associated with changes in gene regulation at several levels (34-37), including: (i) GR induction of IκBα, which binds to the pro-inflammatory transcription factor complex NFκΒ and prevents it from entering the nucleus; (ii) inhibition of histone acetylation at the promoters of pro-inflammatory genes; (iii) post-transcriptional degradation of inflammation-related mRNAs; (iv) recruitment of negative elongation factor at promoters of inflammatory genes, which inhibits activation; and (v) GR tethering to the NFκB component p65 (RelA) at NFκB binding sites to inhibit pro-inflammatory gene transcription. Glucocorticoid resistance, which impairs termination of pro-inflammatory signaling, is thus also linked to multiple regulatory mechanisms, including defective histone acetylation, increased expression of pro-inflammatory transcription factors, and disinhibition of pro-inflammatory signaling by the alternative GR receptor, GRβ (reviewed in (38)).

HPA axis signaling via GC regulation therefore provides a plausible mechanism for linking the central nervous system, which senses and processes social experiences, to gene regulation in the periphery, where social stress is translated into poor health outcomes. Indeed, observational studies of human subjects (39-45) and experimental animal models for social subordination-induced stress (8, 9) have identified a strong relationship between the chronic social stress associated with low social status and pro-inflammatory gene expression in peripheral immune cells. In support of a role for GC signaling, predicted GR and NFκB binding sites are enriched near genes that are differentially expressed in association with social isolation (46), low social status (8, 44, 47), and early life social adversity (48, 49). In addition, experimental manipulation of social status in rhesus macaques results in a more pronounced pro-inflammatory gene expression response to lipopolysaccharide exposure in low-ranking animals. Notably, transcription factor binding sites for NFκB are enriched in accessible chromatin near genes affected by social status, suggesting that changes in GC-related NFκB transcriptional activity may explain the low status-driven pro-inflammatory phenotype (8). However, although low status macaques exhibit some evidence of GC resistance (17-19), both the degree to which GC signaling is responsible for social status-driven gene expression patterns and the gene regulatory mechanisms involved remain unclear.

To address these gaps, here we combined a powerful nonhuman primate model for investigating the consequences of chronic social stress (8, 9) with a well-established model for investigating the gene regulatory response to GC signaling: *in vitro* treatment with the synthetic GC Dexamethasone (36, 50, 51). Specifically, we sequentially introduced 45 female rhesus macaques into nine new social groups of five females each, a manipulation that produces a strong relationship between introduction order and dominance rank (earlier introduction predicts higher social status/dominance rank: (9, 52)). This approach allows the causal effects of social subordination-induced stress to be disambiguated from alternative explanations, while taking advantage of the natural tendency of female rhesus macaques to form stable status hierarchies (53). To test how social status affects the response to GC signaling, we then cultured peripheral blood mononuclear cells (PBMCs) from each study subject using a paired control (untreated) and Dexamethasone-treated study design (following (50)).

Because the majority of GR-DNA binding events are thought to occur in regions of chromatin accessible to TF binding prior to GR nuclear translocation (54), we measured both genome-wide gene expression and chromatin accessibility in our samples. We hypothesized that social status would not only predict variation in PBMC gene expression (as previously shown in macaques and humans (9, 44)) but also patterns of open chromatin. Further, as shown for exposure to LPS (8), we hypothesized that the effects of social status on gene regulation would be conditional on GC exposure. Finally, we hypothesized that low-ranking animals, who exhibit signatures of social stress-mediated GC resistance (23, 29), would also exhibit gene regulatory patterns consistent with reduced responsiveness to GC signaling, particularly at GR and NFκB-regulated genes.

## RESULTS

### Social status and GC stimulation alter chromatin accessibility and gene expression

We manipulated the dominance rank positions of 45 adult female rhesus macaques by sequentially introducing them into newly constructed social groups of 5 females each (as reported in (8)) (Fig. 1A; Table S1). This approach produces a strong correlation between order of introduction and resulting dominance rank, in which females introduced earlier become higher ranking (Pearson’s r = −0.67, p=8.8×10^−7^ at time of sample collection; rank was measured using a continuous Elo rating score; (55); Fig. 1A) (see also (8, 52)). Consistent with dominance rank dynamics typical for female rhesus macaques, dominance rank values were highly stable from group construction until sample collection for the present study.

**Figure 1.**
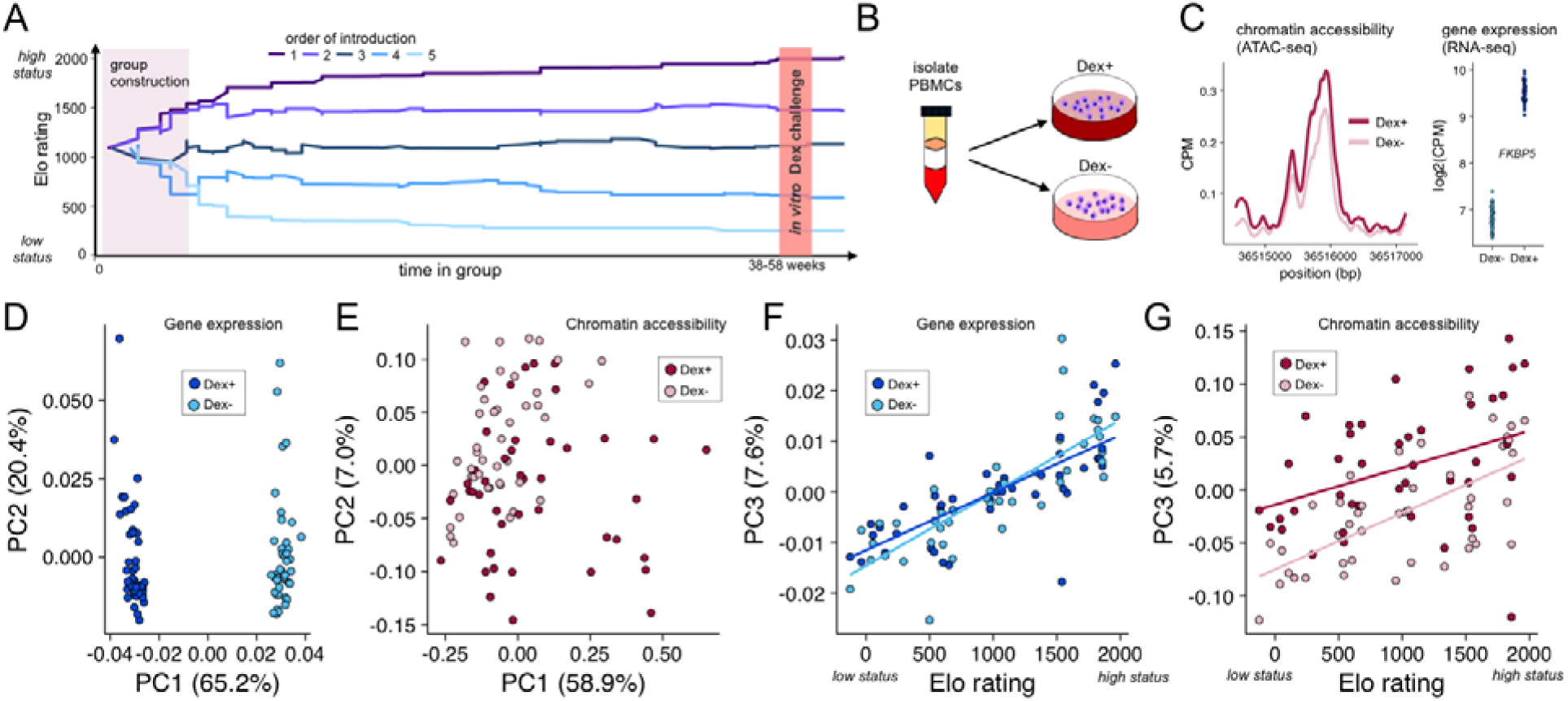
Dominance rank and Dexamethasone treatment both induce widespread changes in gene expression and chromatin accessibility. (A) Social group formation for one study group. Each line represents a different female and starts at the date of her introduction. Females introduced earlier were higher ranking at the time of sampling (Pearson’s r=-0.67, p=8.8×10^−7^; n=9 groups and 43 females). (B) Schematic of the *in vitro* GC challenge. Paired peripheral blood mononuclear cell (PBMC) samples from each female were treated with either 1 µM Dex (“Dex+”), a synthetic glucocorticoid, or 0.02% ethanol (the vehicle for Dex; “Dex-”), and subjected to a 90-minute incubation period before RNA and DNA extraction. (C) Example response to Dex challenge for *FKBP5*, a known GC-induced gene, for chromatin accessibility in intron 4 of isoform *FKBP5-201* (p=7.36×10^−18^; this region also falls in intron 5 of isoform *FKBP5-202*) and for gene expression (p<x10^−100^). (D) Dex treatment is strongly associated with the first principal component of gene expression data (paired T_42_=80.61, p=1.12×10^−47^) and (E) with the first three principal components of chromatin accessibility (PC1 T_42_=-3.85, p=3.95×10^−4^; PC2 T_42_=4.49, p=5.44×10^−4^; PC3 T_42_=-4.93, p=1.35×10^−5^). Social status is significantly associated with PC3 for both (F) gene expression (Pearson’s r=0.77, p=3.33×10^−18^) and (G) chromatin accessibility (Pearson’s r=0.48, p=3.41×10^−6^).

We isolated PBMCs from 43 females (all members of 7 social groups, and 4 of the 5 members of the remaining 2 groups; see Methods). We divided each sample into two aliquots of 1 million cells each, and simultaneously treated one aliquot with 1 µM Dexamethasone (the “Dex+” sample) and the second aliquot with 0.02% EtOH, the vehicle for Dex (the “Dex-“ control sample). Following a 90-minute incubation at 37º C (an incubation period that results in a strong gene expression response to Dex: (50)), we extracted RNA and intact nuclei from each sample for RNA-seq and ATAC-seq analysis, respectively (56, 57). After sequencing and quality control, we retained all 43 control and 43 Dex+ samples for both gene expression and chromatin accessibility analyses (Fig. 1B-C).

As expected (50), Dex treatment resulted in substantial changes to PBMC gene expression levels. Principal components analysis of all expressed genes (n=10,385) clearly separated Dex- from Dex+ samples on PC1 (paired T_42_=80.61, p=1.12×10^−47^; Fig. 1D). At the individual gene level, we identified 7,359 differentially expressed genes (70.9% of those tested) between Dex+ and Dex- samples at a false-discovery rate (FDR) of 10% (n=5,032 differentially expressed genes at FDR<1%) controlling for dominance rank, PBMC composition, and relatedness among study subjects. This set of 7,359 genes (Table S2) included major targets of Dex induction (e.g., *NFKBIA*, *FKBP5*, and *PER1*; Fig. 1C) and repression (*CSF1*, *IER2*, and *NR3C1*, the gene encoding GR) (50) and recapitulated the effect of Dex for Dex-responsive genes identified in previous work in humans or human-derived cell lines (Pearson’s r=0.56, p=3.81×10^−11^, n=119 genes from the A549 lung epithelial cell line (50); Pearson’s r=0.27, p=6.62×10^−25^, n=1450 genes from PBMCs (58); and Pearson’s r=0.40, p=7.19×10^−125^, n=3,232 genes from PBMCs (59)).

Dex treatment also remodeled the PBMC chromatin landscape, although with less pronounced effects than on gene expression. Control and Dex+ samples separated on PCs 1, 2, and 3 of the complete ATAC-seq data set (PC1 T_42_=-3.85, p=3.95×10^−4^; PC2 T_42_=4.49, p=5.44×10^−4^; PC3 T_42_=-4.93, p=1.35×10^−5^; Fig. 1E). Of the 19,859 300-bp windows we analyzed (see Methods), we identified 1,480 (7.45%; full list in Table S3) that became significantly more accessible in the Dex+ condition and 1,602 (8.07%) that become significantly less accessible after Dex treatment, at an FDR threshold of 10% (215 and 190 at an FDR of 1%). Further, although GR is thought to primarily bind DNA in regions that are already accessible prior to nuclear translocation (54), regions with increased accessibility after Dex administration were significantly enriched for two predicted GR binding motifs (log_2_odds=1.75 and 2.04, p=1.86×10^−16^ and 9.56×10^−30^) and for binding sites for the GR co-factor activator protein 1 (Jun-AP1 complex; log_2_odds =1.45, p=3.45×10^−23^; Table S3). Conversely, Dex-repressed regions were enriched for TF binding sites for a component of the pro-inflammatory transcription factor complex, NF-κB (the NF-κB-p50 binding site: log_2_enrichment=1.20, p=1.57×10^−15^; Table S4).

Social status also exhibited a detectable relationship with PBMC gene regulation, although these effects were modest compared to the effects of Dex treatment. In both the gene expression and chromatin accessibility data, dominance rank was strongly associated with sample loading on PC3 (gene expression: Pearson’s r=0.77, p=3.33×10^−18^, Fig. 1F; chromatin accessibility: Pearson’s r=0.48, p=3.41×10^−6^, Fig. 1G). For the gene expression data, the effects of social status for individual genes in the Dex- condition were broadly correlated with those described in previous work on PBMCs in rhesus macaques (Pearson’s r=0.39, p=4.08×10^−24^; (9)). However, the effects of rank on gene expression levels were substantially weaker than those described for either purified individual cell types or for all pooled white blood cells (including granulocytes) in the same study subjects ((8); Figure S1), potentially because here we performed more sample processing after a longer time delay following blood sample collection than for previous mixed cell populations (8, 9). Thus, at a FDR of 10%, we identified only 69 social rank-associated genes in the Dex- samples (n=1,162 genes at an FDR of 20%). We were also able to detect the first evidence for an effect of social status on locus-specific chromatin accessibility. We identified 159 rank-associated chromatin accessibility windows in the Dex- samples (0.8% of those tested at an FDR<10%; 1,084 at an FDR<20%), controlling for kinship and tissue composition.

### GC treatment-dependent effects of social status

Previous work using LPS stimulation indicates that the effects of social status on gene regulation depend on the cellular environment (8). Specifically, in the same rhesus macaque model used here, LPS stimulation magnifies the effects of social status on gene expression levels: social status-associated genes are more detectable, and the strength of these associations is greater, following LPS exposure. In contrast, our results show that Dex treatment dampens the social status-gene expression relationship. Among genes that were nominally associated with status in either the Dex+ or Dex- conditions (p<0.05; n=2,279 genes), social status effects on gene expression were, on average, 4x stronger in the Dex- condition compared to Dex+ (paired T_2278_=17.33, p=2.63×10^−63^; Fig. 2A). Additionally, with increasing effect size, the number of social status-associated genes detected in the Dex- versus the Dex+ condition became increasingly more biased towards the Dex- condition (Fig. 2B). As a result, there were almost twice as many social status-associated genes in the Dex- versus the Dex+ condition at any p-value threshold below 0.05 (Fig. 2B). This observation is not explained by increased noise in the Dex+ condition (Fig. S2A). However, while social status effect sizes were smaller after Dex treatment, they were generally well correlated between conditions (Pearson’s r=0.75; p<10^−300^; Fig S3). Thus, although the change in effect sizes between Dex- and Dex+ conditions implies an interaction between social status and GC exposure, we did not identify individual genes for which the log-fold change gene expression response to Dex was significantly associated with social status (10% FDR).

**Figure 2.**
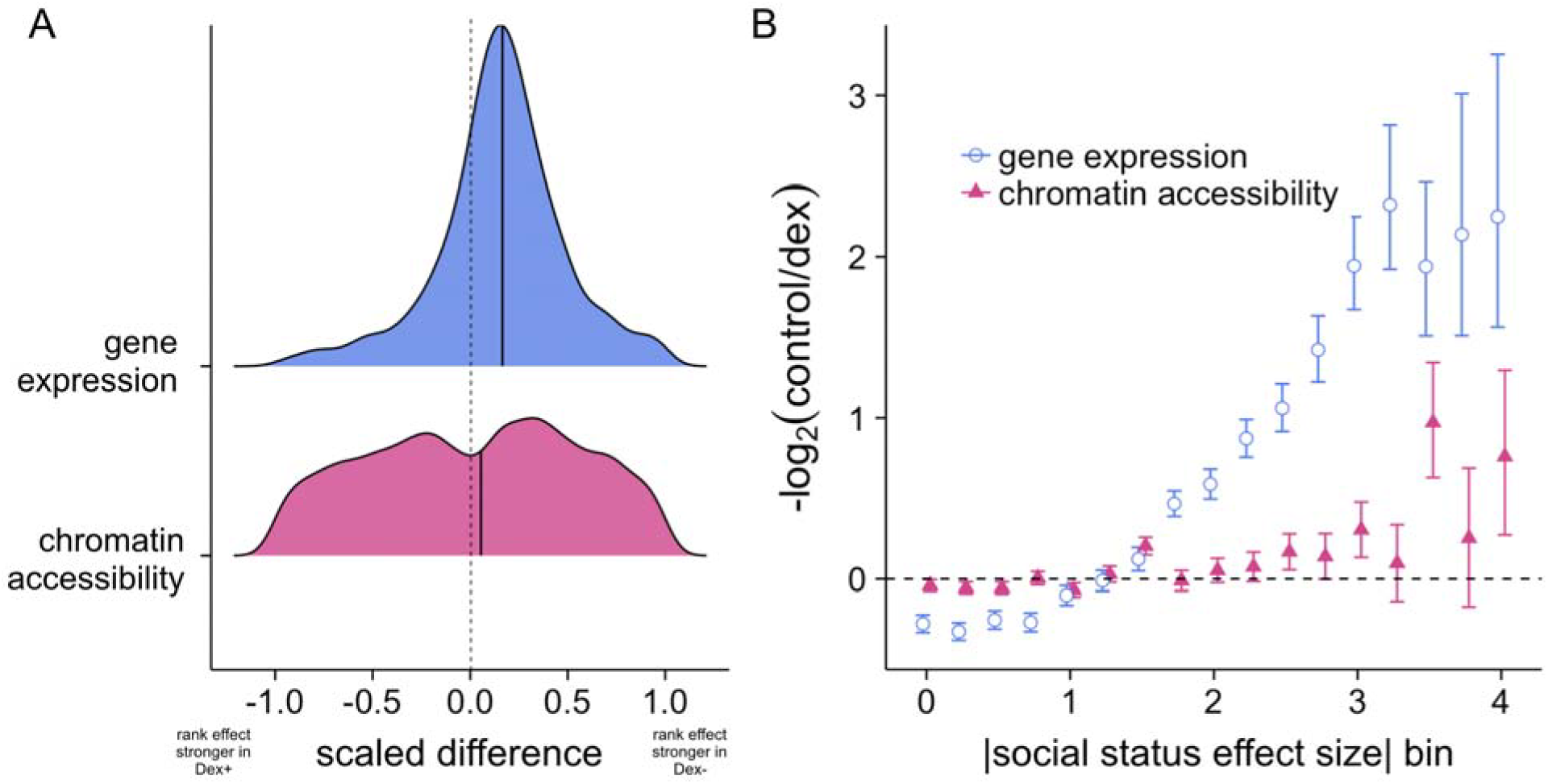
Differences in the stability of social status effects on gene expression and chromatin accessibility after Dexamethasone treatment. (A) The effects of social status on gene expression are stronger before Dex treatment (paired T_2278_=17.33, p=2.63×10^−63^), but status effects on chromatin accessibility are similar before and after Dex treatment (paired T_4082_ =2.69, p=0.007). “Scaled difference” is the difference between the absolute value of the effect of social status in the Dex- and Dex+ conditions scaled by the sum of the effect sizes: 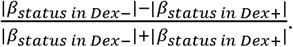 Solid vertical lines represent the median difference in scaled effect sizes. (B) We observed an excess of genes with large social status effects on gene expression in the Dex- versus the Dex+ condition. In contrast, we observed no consistent bias in the number of social status-associated genes in the Dex-versus Dex+ condition as effect size increased. The x-axis depicts mutually exclusive bins of effect sizes for the standardized effects of status on gene expression (blue) or chromatin accessibility (pink). The y-axis represents the log_2_-transformed ratio (± se) of the number of genes falling in each bin for the Dex- compared to the Dex+ condition.

Compared to social status effects on gene expression, social status effects on chromatin accessibility were stable between conditions. In regions nominally associated with rank in either condition (p<0.05; n=4,083 regions), the effect of social status was significantly different across Dex- and Dex+ conditions, but much more similar in direction and magnitude than for gene expression (paired T_4082_=2.69, p=0.007; Fig. 2A). Additionally, unlike for gene expression, we observed no evidence for stronger rank-chromatin accessibility associations in the Dex- relative to the Dex+ condition with increasing effect size (Fig. 2B), despite higher variance in the Dex+ condition (Fig. S2B). Thus, while GC treatment rapidly altered social status effects on gene expression levels, it did not induce systematic changes in social status-associated chromatin accessibility on the same timescale.

### Dex treatment decouples social status-associated chromatin regions from social status-associated gene expression

To test whether social status effects on gene expression are explained by social status-driven differences in chromatin accessibility, we next examined the relationship between the RNA-seq and ATAC-seq data sets. In the Dex- condition, we found that social status effects on chromatin accessibility significantly predicted the magnitude and direction of social status effects on the expression level of the closest gene (Pearson’s r=0.23, p=1.51×10^−199^; n=16,677 region-gene pairs; Fig. 3A). Specifically, regions that were more accessible in high-ranking females were associated with genes that were also more highly expressed in high-ranking females (and vice-versa). This relationship becomes stronger as either the threshold for classifying social status-associated genes (Fig. 3C) or the threshold for classifying social status-associated chromatin accessibility regions becomes more stringent (Fig. 3D). Thus, while social status effects on chromatin accessibility explained only 5.3% (95% CI: 4.6-6.0%) of variance in social status effects on gene expression for all region-gene pairs, they explained 12.7% (95% CI: 10.7-14.9%) of variance of social status-driven gene expression for genes where rank was at nominally associated with gene expression (p < 0.05). This estimate is almost certainly conservative, as the regulatory elements that influence a gene’s expression are not well-predicted by proximity alone (60). Thus, under baseline (e.g., non-GC stimulated) conditions, social environment-driven changes in the chromatin landscape may mechanistically contribute to social environment-driven variation in gene expression levels.

**Figure 3.**
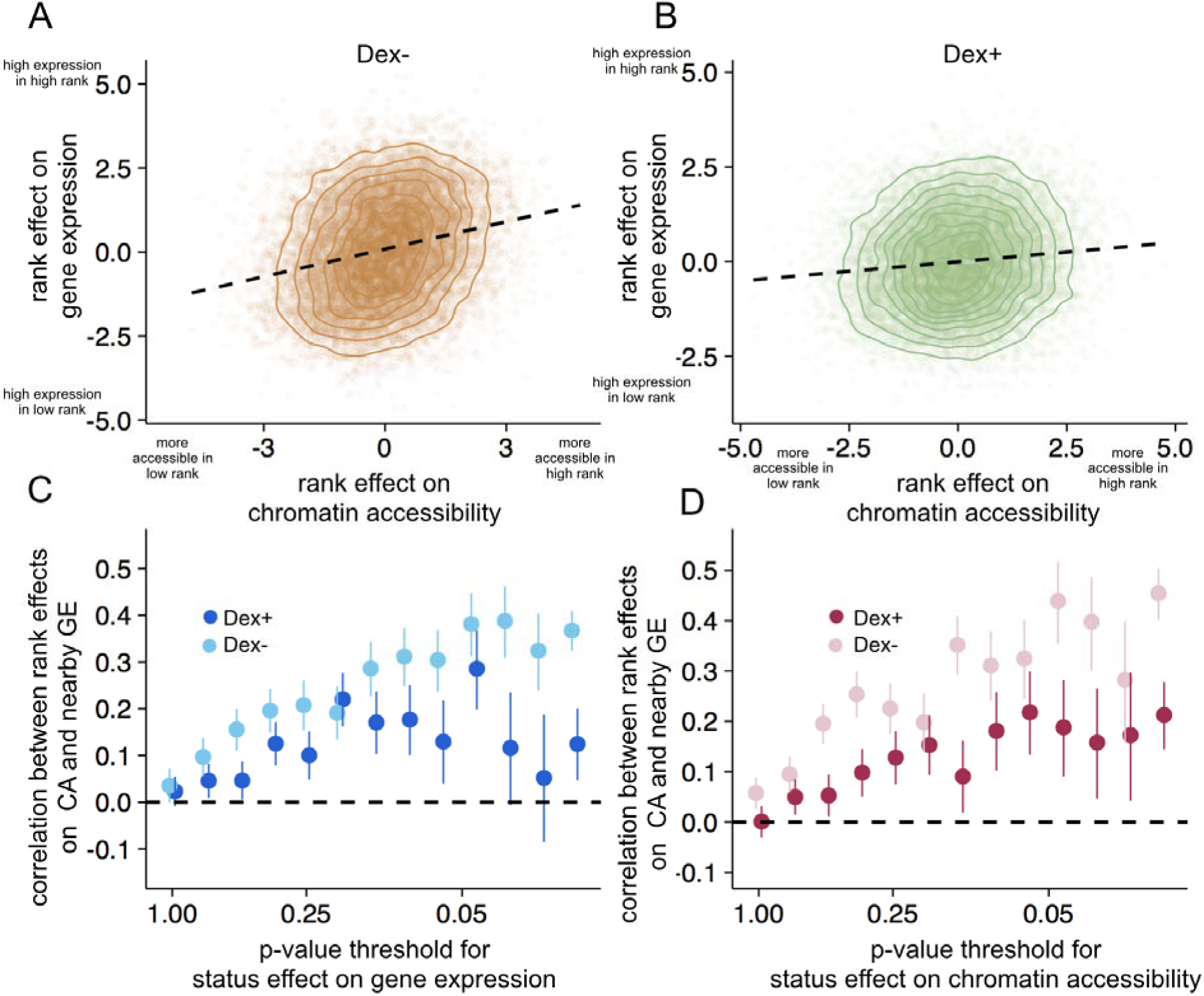
Dex treatment affects the correlation between rank-associated chromatin accessibility and rank-associated gene expression levels. Standardized rank effects on chromatin accessibility and standardized rank effects on the expression of the closest gene are correlated in the Dex- condition (Pearson’s r=0.23, p=1.51×10^−199^; n=16,677 region-gene pairs) (A) but negligibly so after Dex treatment (r=0.10, p=2.76×10^−9^; n=16,677) (B). (C,D) Pearson’s correlation (± se) between the standardized effect of social status on chromatin accessibility and the standardized effect of social status on gene expression for the closest gene (y-axis) is stronger in the Dex- condition compared to the Dex+ condition, whether conditioning on the gene expression association with dominance rank (C, x-axis) or the chromatin accessibility association with dominance rank (D, x-axis). CA: chromatin accessibility; GE: gene expression.

Social status effects on chromatin accessibility and social status effects on the expression levels of nearby genes were also significantly correlated after Dex stimulation (r=0.10, p=2.76×10^−9^; n=16,677 region-gene pairs; Fig. 3B). However, the magnitude of this correlation was reduced more than 2-fold relative to before treatment (Dex-). Social status effects on chromatin accessibility explained only 1.07% of the variance in social status effects on gene expression in the Dex+ data, and this relationship was not appreciably strengthened by altering the statistical threshold for identifying status-associated genes or regions (Fig. 3C & D, dark colors). These results indicate that social status effects on different levels of gene regulation are partially decoupled following environmental challenge with glucocorticoids, at least on a short-term timescale.

### Shared chromatin accessibility signatures of social status and GC treatment

Previous studies in female rhesus macaques have identified evidence for higher GC levels and increased GC resistance in low status animals (reviewed in (61)), and chronic social stress effects on immune gene expression have been hypothesized to originate in part from GC dysregulation (23). If social status effects operate through a GC-mediated pathway, then gene regulatory phenotypes affected by social status should overlap with those affected by Dex treatment.

This prediction was better supported by the chromatin accessibility data than the gene expression data. The effects of rank on gene expression in unchallenged cells and the effects of Dex on gene expression were not correlated (Pearson’s r=0.007, p=0.47; Fig. 4A). Further, rank effects were actually stronger in Dex-unresponsive genes (Dex effect FDR>20%) than Dex-responsive genes (Dex effect FDR<1%; T_2014_=5.10, p=3.66×10^−7^). In contrast, rank effects on chromatin accessibility in unchallenged cells were much more strongly correlated with Dex effects on chromatin accessibility, such that regions that were more accessible after Dex treatment were also more accessible in high-ranking females, and vice versa (Pearson’s r=0.18, p=5.78×10^−147^; Fig. 4B). Rank-associated chromatin accessibility windows and Dex-associated chromatin accessibility windows were also functionally linked through their component transcription factor binding site (TFBS) motifs. Specifically, TFBS motifs enriched in regions that were more accessible in high-ranking individuals were similarly enriched in regions that opened after Dex treatment (Pearson’s r=0.64, p=1.95×10^−41^; Fig. 4C; Table S4). Compared to PBMCs from low status individuals, cells from high status individuals thus appear epigenetically primed to respond to glucocorticoids. As was the case with gene expression, we found little evidence for rank by treatment interactions on chromatin accessibility: rank predicted the log_2-_foldchange chromatin accessibility response to Dex at 3 loci (FDR<10%; n=31 loci at an FDR<20%).

**Figure 4.**
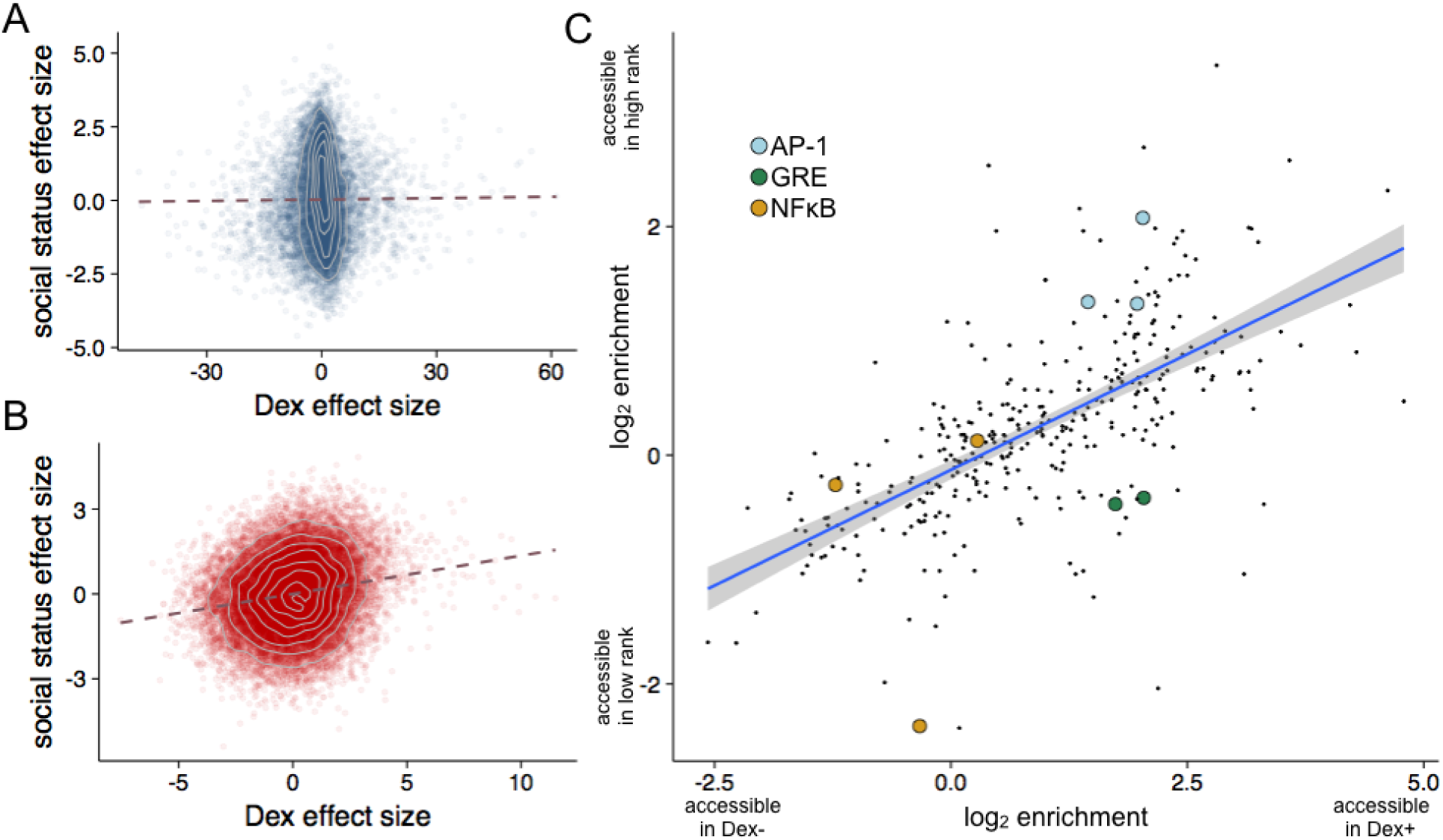
The relationship between rank effects and Dex effects on gene expression, chromatin accessibility, and TF accessibility. (A) The effects of rank and Dex treatment on gene expression were uncorrelated (Pearson’s r=0.007, p=0.47) (B) the effects of rank and Dex treatment on chromatin accessibility (Pearson’s r=0.18, p=5.78×10^−147^). (C) Transcription factor (TF) binding sites enriched in Dex-induced regions (n=2,874 regions, FDR<20%) also were enriched in regions that were more accessible in high-ranking females (n=544 regions, FDR<20%; Pearson’s r=0.64, p=1.95×10^−41^). Conversely, TF binding sites enriched in Dex-repressed regions (negative values on the x-axis; n=2,905 regions, FDR<20%) were also enriched in regions that were more accessible in low-ranking females (negative values on the y-axis; n=540 regions, FDR<20%) Each point (n=369 TFBS) represents the log_2_-foldenrichment of one TF motif in Dex-induced compared to Dex-repressed regions (x-axis) and rank-induced vs. rank-repressed regions (y-axis). TF motifs closely associated with GC regulation (GRE, AP-1, NFκB) are shown in colored circles.

## DISCUSSION

Together, our results reinforce both the importance of social interactions for immune cell gene regulation and the context-dependent nature of this relationship. Specifically, we provide the first support for systematic, status-induced changes in chromatin accessibility in primary immune cells, adding to the growing literature on social status effects on gene expression (8, 9, 39, 42, 44, 47, 62, 63), DNA methylation (9, 64-67), and H3K27ac (68). Rank-driven variation in chromatin accessibility may also contribute to previously reported rank-driven variation in the gene expression response to lipopolysaccharide (8). Our results show that low ranking animals present chromatin landscapes more accessible at some TFBSs for NFκB, a key transcription factor in the inflammatory response. In contrast, high ranking females exhibit more accessible binding sites for the GR co-factor AP-1, which is involved in anti-inflammatory activity and NFκB repression (Fig. 4C).

While social status effects are exaggerated after exposure to LPS (8), here we found that social status effects are attenuated after glucocorticoid treatment (Fig. 2A). Our results are broadly consistent with the evidence that chronic social stress leads to poor control of the inflammatory response (23). Specifically, by administering a high dosage of anti-inflammatory glucocorticoids, we appear to have attenuated, and in some cases eliminated, the signature of low social status. However, we note that Dex treatment did not entirely eliminate the gene regulatory signature of social status. High-ranking and low-ranking females presented consistently different patterns of chromatin accessibility regardless of treatment condition, resulting in a detectable association between social status effects on gene expression and chromatin accessibility at baseline, but not following Dex administration. These observations suggest that the imprint of chronic stress exposure may persist in the epigenome even when gene expression levels shift in response to short-term environmental cues. Such a model is appealing because the effects of social adversity are sometimes observable over the span of years, suggesting that they remain “biologically embedded” even as individuals experience and respond to other environmental stimuli (69). Some degree of decoupling between the gene regulatory substrates for biological embedding and gene expression levels themselves could help satisfy this requirement for simultaneous stability and plasticity.

A non-mutually exclusive alternative explanation is that we stopped the experiment too early to observe treatment effects on social status-associated chromatin accessibility comparable to those observed for gene expression. We chose our sample collection time because it yields a strong gene expression response to Dex (50), consistent with the idea that changes in chromatin accessibility facilitate downstream changes in gene expression. However, recent evidence suggests that gene expression changes occur rapidly in response to a stimulus and often precede detectable changes in the epigenome (e.g., CpG methylation: (70, 71)). After stimulation by glucocorticoids, many GR-DNA binding events occur at sites that are already accessible prior to GC stimulation (32). Thus, changes in social status effects on chromatin accessibility might not manifest until a later time point (e.g., as a consequence of the secondary effects of GC signaling and GR activation, as opposed to the primary response). Such a possibility would be consistent with our observation that the gene expression data are more clearly separated by treatment condition than the chromatin accessibility data after 90 minutes (Fig. 1D-E). Future studies with more time points would enable finer dissection of social status effects on the gene regulatory response to GCs, including whether social status affects how quickly cells return to a baseline state.

Our findings combine with those of others (8, 59, 70, 72) to emphasize that the detectability and magnitude of predictors of gene expression variation are often context-dependent. This idea has attracted substantial interest in research on the genetic contribution to gene regulation, where an increasing number of studies have identified genetic variants that influence the *response* to environmental stimuli (i.e., “response-QTL”) as opposed to, or in addition to, gene expression levels at baseline (e.g., (59, 73-78)). Recent work suggests that response-QTL can emerge as a consequence of genetic variation that influences chromatin accessibility in enhancer regions prior to stimulation (79). Under this “enhancer priming” model, changes in the cellular environment post-stimulation alter the complement of active transcription factors, thus translating previously silent effects of genotype on chromatin architecture to detectable effects on gene expression. Our results suggest that social environmental effects on gene expression can be masked or unmasked by environmental stimuli in a similar fashion. Here, status-driven differentially accessible chromatin predicts status-driven differential gene expression under some conditions (Dex-) but not others (Dex+), likely in relationship to the activity of GC-associated transcription factors.

Taken as a whole, our results indicate a subtle, but potentially important role for GC physiology in mediating the effects of social status on PBMC function. Key aspects of PBMC chromatin accessibility in high status females—particularly increased accessibility to the GR co-factor AP-1 and decreased accessibility to NFκB—overlap with Dex-induced changes. Given that GR binding is largely predicted by the chromatin accessibility landscape prior to GC stimulation (54, 80, 81), we speculate that increased accessibility at GR co-factor binding sites in high ranking animals at baseline could drive more efficient GC negative feedback (82), which in turn would result in tighter control of the inflammatory response. If so, increasing chromatin accessibility at key glucocorticoid-sensitive targets (or decreasing accessibility at key NFκB targets) in cells from low-ranking animals (for example, by using epigenome editing approaches (83)) would be expected to reconcile some of the gene expression differences between low- and high-status individuals. Notably, our results also suggest that, while most research on the gene expression response to GC stimulation has focused on its genetic determinants (51, 84, 85), social environmental factors may also be an important factor to consider in human population samples.

## METHODS

### Study population

Samples were collected from 43 captive female rhesus macaques (*Macaca mulatta*) at the Yerkes National Primate Research Center in March 2015. These study subjects were part of a multi-year study on the effect of dominance rank (the primary measure of social status in rhesus macaques) on immune function. Demographic details for each study subject are provided in Table S1. The study commenced in January – June 2013 using 45 females (5 females assigned to each of 9 social groups), as described in (8). In March – June 2014, social group membership was rearranged to form 9 new groups in which females that occupied the same or adjacent ordinal ranks in the initial social groups were co-housed together. This approach maximized all possible changes in dominance rank between the initial and rearranged groups, a strategy that allows us to establish the causal relationship between dominance rank and downstream outcomes. Social groups were constructed by sequentially introducing females into indoor-outdoor run housing (25 m by 25 m for each area) over the course of 2-15 weeks, and behavioral data collection started after the fifth and final female was introduced into each group.

For the present analysis, we used only samples collected in the second phase of the study (study groups formed in March – June 2014). One of the original study subjects died before we could sample her, and a second female was ill at the time of sampling so we excluded her from the current analysis, leaving a total of 43 study subjects. Samples were collected 38–50 weeks after the fifth and final female was introduced to each social group, after rank hierarchies had stabilized following the group rearrangement. Behavioral data in this study included a mean of 19.5 hours of observational data per group (range=14.5-23.0 hours), collected using focal sampling (86). We used the *EloRatings* package in R to assign social status using the Elo method, a continuous measure of dominance rank that is based on an animal’s history of dominance interactions with its group mates (55, 87, 88). The Elo stability index ranged between 0.995 – 1.00 (n = 9 groups), where 1 represents a hierarchy in which higher-ranking females always win competitive encounters with lower-ranking females, with no rank reversals (89), and 0 represents one in which rank does not predict the outcomes of competitive interactions.

### *In vitro* Dexamethasone treatment

For each study subject, we drew 4 mL of blood into BD Vacutainer CPT™ tubes and shipped the blood sample overnight on ice from the Yerkes National Primate Research Center to Duke University, where PBMCs were purified via gradient centrifugation. To control for variation in cell type, an aliquot of 50,000 PBMCs was analyzed by flow cytometry. We quantified the proportion of 10 different cell types in the sample: classical monocytes (CD14^+^/CD16^−^), CD14^+^ intermediate monocytes (CD14^+^/CD16^+^), CD14^−^ non-classical monocytes (CD14^−^/CD16^+^), helper T cells (CD3^+^/CD4^+^), cytotoxic T cells (CD3^+^/CD8^+^), double positive T cells (CD3^+^/CD4^+^/CD8^+^), CD8^−^ B cells (CD3^−^/CD20^+^/CD8^−^), CD8^+^ B cells (CD3-/CD20^+^/CD8^+^), natural killer T lymphocytes (CD3^+^/CD16^+^), and natural killer cells (CD3-/CD16^+^). Cell type composition was summarized using principal components analysis, where the first three PCs of cell composition together explained 97.7% of the variance in whole blood composition. We did not obtain flow cytometry data for two animals, so imputed the values of PCs 1 – 3 for these two samples using elastic net regression fit to the Dex- gene expression data (90).

For each study subject, we seeded at 1 million PBMCs each into two wells of a tissue culture plate containing 2 mL of media (RPMI; 15% FBS; 1% penicillin-streptomycin). We treated one of the wells with 1 µM Dexamethasone (“Dex+”) and the other with 0.02% EtOH (“Dex-”), the vehicle for Dex. Cells were then incubated for 90 minutes (37°C and 5% CO_2_), washed with 1x PBS, and then harvested for gene expression and chromatin accessibility analyses.

### Chromatin accessibility

We measured chromatin accessibility using the ATAC-seq method (56, 57) and 50,000 cells from each treatment-sample combination (n=43 Dex- and 43 matched Dex+). We followed the protocol from (57), but skipped the cell lysis step, a modification that reduces the proportion of reads mapping to the mitochondrial genome. ATAC-seq libraries were amplified for a total of 12 PCR cycles, barcoded, multiplexed, and sequenced on an Illumina NextSeq 500 using paired-end, 38 bp reads. Reads were mapped to the rhesus macaque genome (*Mmul_8.0.1*; Ensembl gene build 87) using the *bwa-mem* algorithm with default settings (91). The resulting data set exhibited the typical periodic pattern of insert sizes associated with nucleosome positioning (Fig. S3). For downstream analyses, we only considered reads that mapped uniquely to the nuclear genome with a mapping quality score ≥ 10. For differential accessibility analysis, we then counted the number of Tn5 insertion sites (5′ of the reads aligned to the positive strand offset by +4 bp and to the negative strands offset by −5 bp) that mapped to non-overlapping 300 bp windows of the genome using the bedtools “coverage” function. Prior to normalization, we filtered all regions that did not pass our coverage threshold (median CPM < 0.5 in the 86 samples) or had poor mappability (<0.90 as measured using GEM (92)).

### Gene expression

Total RNA was extracted from each PBMC sample using Qiagen’s RNeasy Mini kit. RNA-sequencing libraries were prepared from 200 ng of total RNA using the NEBNext Ultra RNA Library Prep Kit, and sequenced on an Illumina NextSeq 500 using paired-end, 38 bp reads. We quantified gene expression levels from the resulting reads by aligning them to the rhesus macaque transcriptome (*Mmul_8.0.1*; Ensembl gene build 87) using *kallisto* (93) and analyzed gene-wise estimates for genes with an RPKM ≥ 2 in at least half the samples from each condition (n=10,385 genes). Details on RNA-seq and ATAC-seq sequencing depth and mapping rates can be found in Table S5.

### Associating genomic regions with the nearest gene

We paired each of the 19,859 non-overlapping windows from the chromatin accessibility analysis to the TSS of the nearest gene. For a subset of these region-gene pairs (n=4,129), the nearest gene was not included in the gene expression data set; these pairs were excluded from downstream analysis. We thus analyzed a total set of 16,677 region-gene pairs, which consisted of 16,102 regions and 7,547 genes.

### Read count normalization and correction for batch effects

We normalized the filtered RNA-seq and ATAC-seq read count matrices using the function *voom* from the R package *limma* (94). Next, we modeled these normalized values as a function of the social group membership of the sample donor (9 total social groups). This allowed us to remove biological variation related to differences in group dynamics and, because we sampled all females in each group on the same day, most technical batch effects related to sample collection and processing. We then used the residuals from these models as our primary outcome measure in all subsequent analyses.

### Modeling rank and treatment effects on gene regulation

We used a linear mixed-effects model that controls for relatedness to identify genes or regions that were significantly associated with Dex treatment or dominance rank (95-97). We analyzed the normalized chromatin accessibility and normalized gene expression data sets separately using the R package EMMREML (98). We estimated the effects of rank and treatment on chromatin accessibility in each region and expression of each gene using the following model:

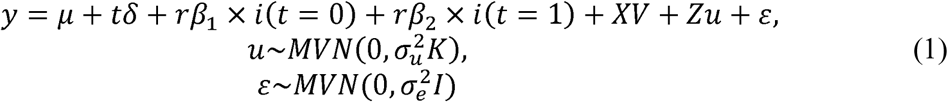

where *y* is the 86 sample vector of normalized gene expression levels or chromatin accessibility levels for a given gene/region; *µ* is the intercept; *t* is the treatment (Dex- = 0, Dex+ = 1); *r* is Elo rating (i.e., social status); and *i* is an indicator variable for the treatment (Dex- = 0, Dex+ = 1); *β_1_* is the effect size of status in the Dex- condition; and *β_2_* is the status effect in the Dex+ condition; and is the effect of Dex treatment. The *m* by 1 vector *u* is a random effects term to control for kinship and other sources of genetic structure. Here, *m* is the number of unique females in the analysis (*m*=43), the *m* by *m* matrix *K* contains estimates of pairwise relatedness derived from a 54,165-variant genotype data set described in (8), 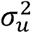 is the genetic variance component (0 for a non-heritable trait; note that most gene expression levels are heritable: (99, 100)), and *Z* is an incidence matrix of 1’s and 0’s that maps measurements in the Dex+ and Dex- conditions to individuals in the random effects term, thus accounting for repeated measurements for the same individual. Residual errors are represented by *ε*, an n by 1 vector, where 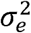 represents the environmental variance component (unstructured by genetic relatedness), *I* is the identity matrix, and MVN denotes the multivariate normal distribution. Other fixed effects covariates included were the first three principal components of the cell type proportion data, which together explain 97.7% of the variance in whole blood composition (matrix *X*). For each data set and gene, we tested the null hypothesis that the effect of interest (*β* or *δ*) was equal to 0 versus the alternative hypothesis that the effect was not equal to 0.

For all models, we produced empirical null distributions for rank effects by permuting the Elo rating values associated with pairs of samples (Dex- and Dex-treated) as a block, 1000 times, and re-running our analyses. To produce a corresponding empirical null for the effect of treatment, we randomized treatment (Dex- versus Dex+) within the pair of samples for each female.

For all of our analyses, we controlled for multiple testing using a generalization of the false discovery rate method of Storey and Tibshirani (101) to empirical null p-value distributions generated via permutation (code available at https://github.com/nsmackler/status_genome_2016/blob/master/perm_FDR.R).

### Transcription factor binding site (TFBS) enrichment

To test if binding sites for specific TFs were consistently found in social status- or Dex-associated chromatin regions, we used the motif analysis program HOMER (102) and the function “findMotfsGenome.pl”. We conducted two comparisons to test for the enrichment of 309 known vertebrate binding motifs: (i) regions that were more open in high status animals compared to regions that were more open in low status animals (FDR < 20%; n=544 regions more open with higher rank, n=540 regions more open with lower rank), and (ii) regions that were more open after Dex treatment compared to regions that were less open after Dex treatment (FDR < 20%; n=2,854 Dex-induced regions, n=2,902 Dex-repressed regions).

## Author contributions

J.T., L.B.B., M.E.W., and N.S-M. designed the research; J.T., N.S-M., J.K, T.V., and L.B.B. performed research; J.T., L.B.B., N.S-M., J.S., and R.P-R. contributed new reagents/analytic tools; N.S-M., J.S., J.T., L.B.B., and R.P-R. analyzed data; and N.S-M., J.T., L.B.B., and J.S. wrote the paper with contributions from all coauthors.

## Acknowledgements

We thank J. Whitley, A. Tripp, N. Brutto, and J. Johnson for maintaining the study subjects and collecting behavioral data; A. Lea and C. Vockley for thoughtful comments on the manuscript; and members of the Tung lab for helpful discussion. This work was supported by NIH grants R01-GM102562, P51-OD011132, K99/R00-AG051764, and T32-AG000139; NSF grant SMA-1306134; the Canada Research Chairs Program 950-228993; and NSERC RGPIN/435917-2013. J.S. was supported by the Fonds de recherche du Québec–Nature et technologies, the Fonds de recherche du Québec–Santé, and the Canadian institutes of Health Research Banting fellowship. RNA-seq and ATAC-seq data have been deposited in Gene Expression Omnibus (SRP150653). Code and data are available at github.com/nsmackler/Dex_status_2018.

## Supplemental Figures

**Supplemental Figure 1.**
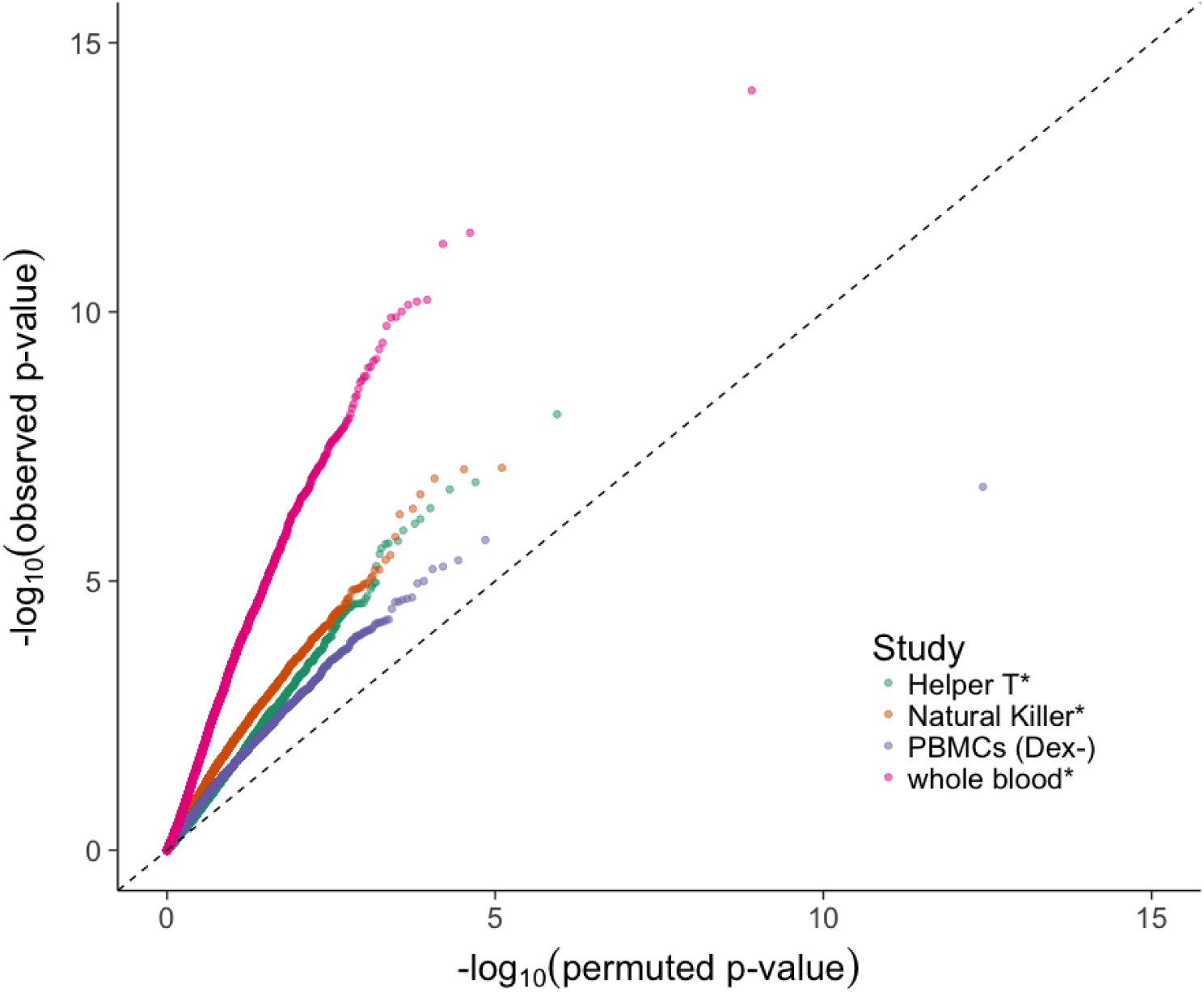
Quantile-quantile plot of the effects of dominance rank on gene expression from this study and Snyder-Mackler et al (2016). The PBMC samples used in this analysis (purple) underwent a longer delay until processing relative to the whole blood samples, and contained a more heterogeneous mixture of cells than the NK and helper T cell samples. We speculate that these differences explain lower detectability of social status-gene expression associations here compared to previous work.

**Supplemental Figure 2.**
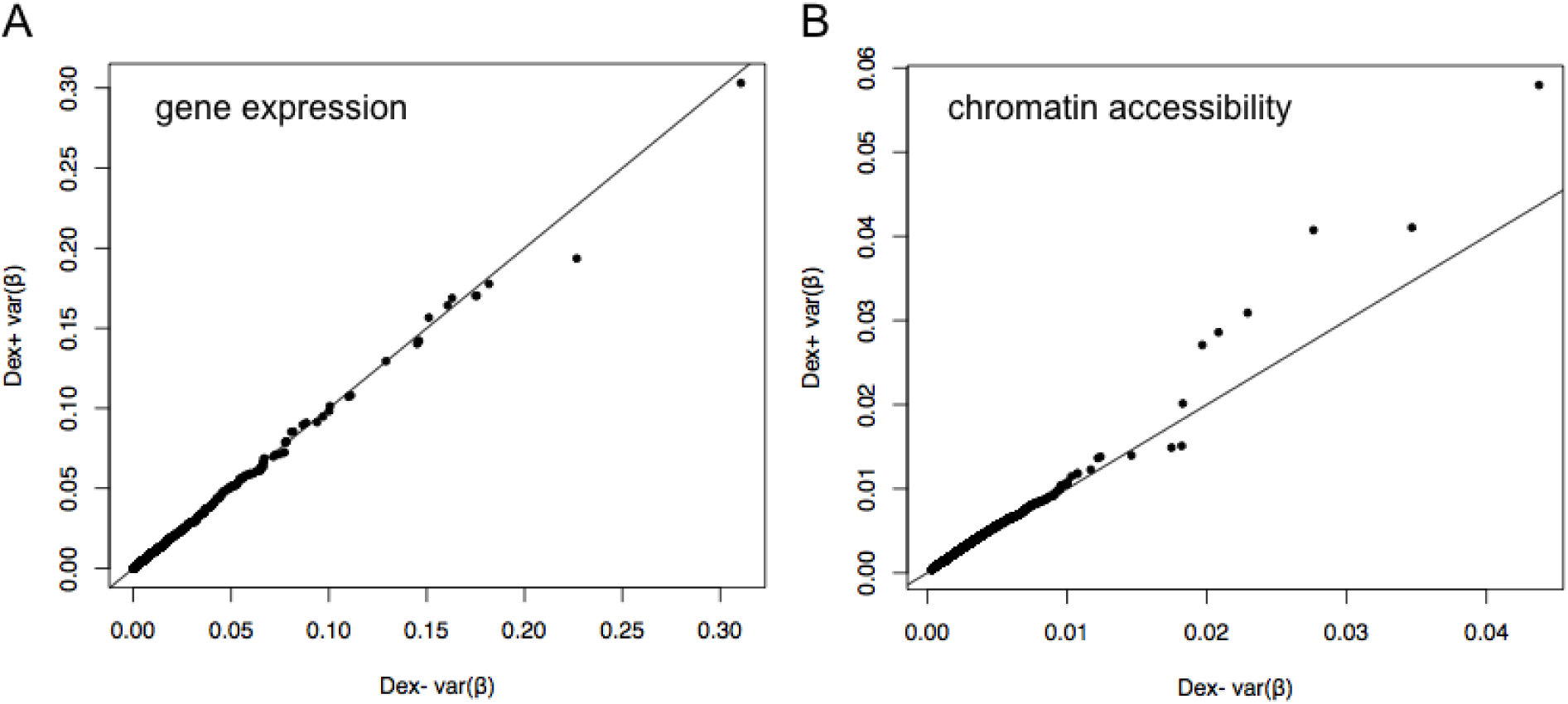
(A) We observed no significant difference in the estimates of variance in rank effects on gene expression in the Dex- condition compared to the Dex+ condition (paired T=1.24, p=0.21). (B) However, our estimates of variance in rank effects on chromatin accessibility were significantly larger after Dex treatment (paired T=51.283, p<10^−300^).

**Supplemental Figure 3.**
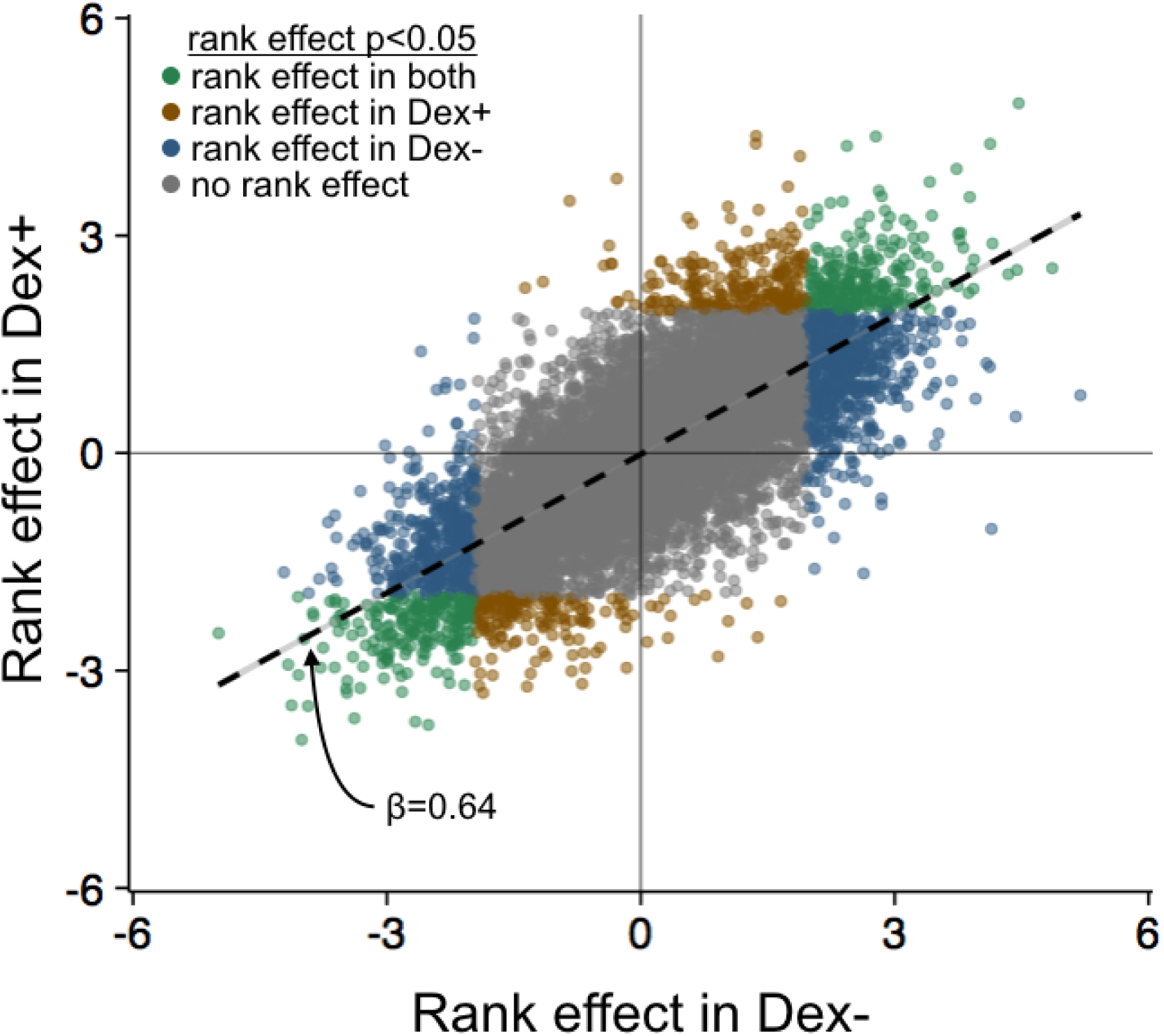
Correlation between the standardized effect of social status on gene expression in the Dex- and Dex+ conditions (r=0.77; p<10^−300^). 97% (2,205/2,279) of the genes that were significantly associated with rank at p < 0.05 in either condition were concordantly affected by rank. However, the effect of rank on gene expression was significantly stronger in the Dex- condition compared to the Dex+ condition.

**Supplemental Figure 4.**
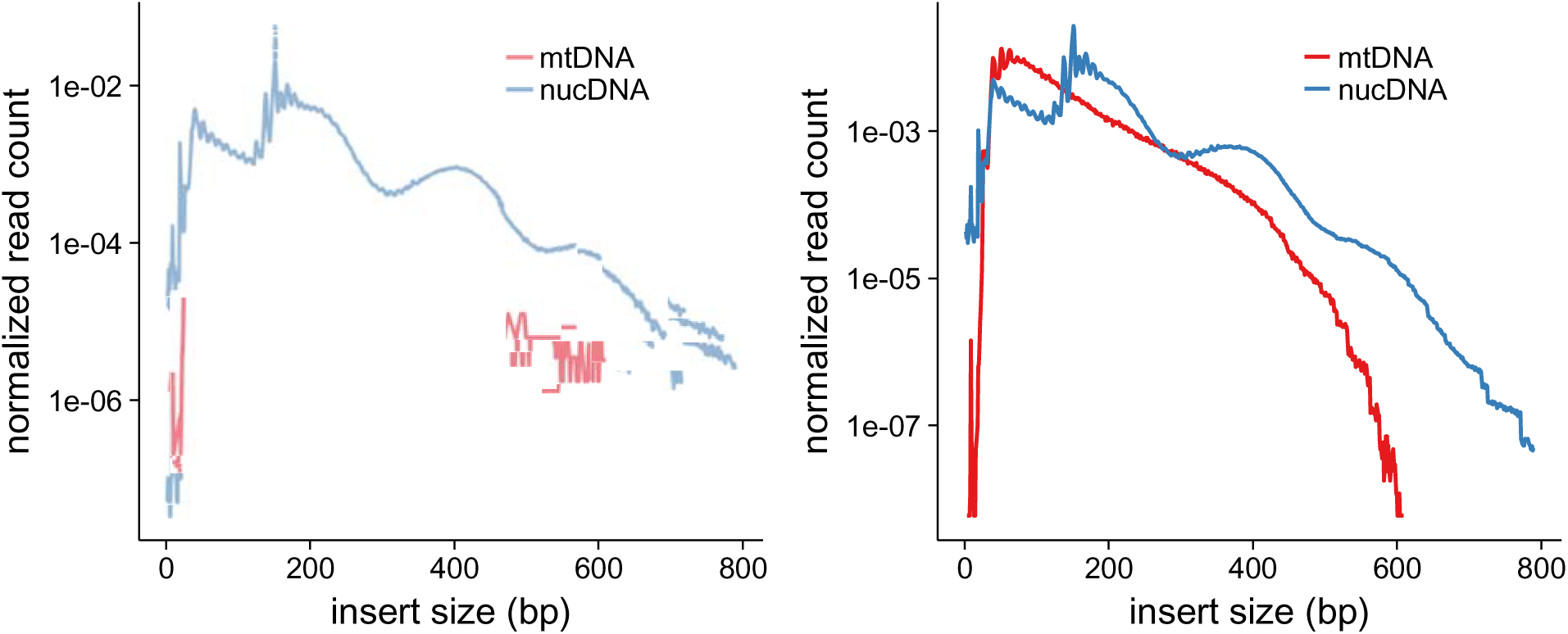
ATAC-seq library insert size distribution (per sample on the left and combined across all samples on the right)

